# Application and comparison of lyophilisation protocols to enhance stable long-term storage of filovirus pseudotypes for use in antibody neutralisation tests

**DOI:** 10.1101/2022.05.13.491820

**Authors:** Martin Mayora Neto, Edward Wright, Nigel Temperton, Peter Soema, Rimko ten Have, Ivo Ploemen, Simon Scott

**Affiliations:** Viral Pseudotype Unit (VPU), Medway School of Pharmacy, Universities of Kent and Greenwich at Medway, Central Avenue, Chatham Maritime, ME4 4TB; Viral Pseudotype Unit, University of Sussex, Falmer, Brighton BN1 9RH; Intravacc, P.O. Box 450, 3720 AL, Bilthoven, The Netherlands

## Abstract

Filoviruses encompass highly pathogenic viruses placing sporadic public health burden on countries affected. Efforts for improved diagnostics and surveillance are needed considering the recent Ebola outbreaks in Africa. The need for high containment facilities can be circumvented by the use of pseudotype viruses (PV), which can be handled in low containment, for tropism, drug screening, vaccine immunogenicity and serosurveillance studies.

In this study we assessed stability and functionality after long-term storage of lyophilised filovirus pseudotypes for use in neutralisation assays. Lyophilised Ebola and Marburg PVs retained production titres for at least two years when stored at +4°C or less. Lyophilised Ebola PVs performed similarly to non-lyophilised PVs in neutralisation assays after reconstitution. When stored at high temperatures (+37°C), lyophilised PVs did not retain titres after one-month storage, however, when lyophilised using pilot scale facilities EBOV PVs retained titres and performed well in neutralisation assays after one-month storage at 37°C suggesting removing residual moisture might be crucial for avoiding cold-chain transportation. Lyophilisation could allow reagents to be transported more efficiently as well as reducing costs for a future serological kit.

## 1. Introduction

Filoviruses have been responsible for several serious disease outbreaks within resource-limited countries, which have posed challenges for implementation of appropriate public health measures. These sporadic Ebola and Marburg virus outbreaks culminated in the large epidemic in the Democratic Republic of Congo in 2013-2016, highlighting the need for better serosurveillance, diagnostics, containment measures, treatments and vaccines (Languon and Quaye 2019).

The gold standard for diagnostics of filoviruses is viral RNA detection via RT-qPCR, which has high sensitivity and specificity but requires user expertise and expensive equipment (Weidmann, Mühlberger and Hufert 2004; Cherpillod *et al*. 2016; Broadhurst, Brooks and Pollock 2016). Several approaches for point-of-care use are being evaluated. Some of these platforms such as RT-PCR based GeneXpert require minimal training and no sample pre-treatment (Semper *et al*. 2016; Vuren *et al*. 2016; Raftery *et al*. 2018). Portable lateral flow devices for antigen detection are also being evaluated as more affordable options. These exhibit varying degrees of sensitivity, which would have to be addressed before being rolled out (Phan *et al*. 2016; Wonderly *et al*. 2019; Makiala *et al*. 2019). More recently, genomic approaches including next-generation sequencing platforms have been employed for diagnostics, as well as monitoring geographical spread and adaptations as an epidemic progresses (Gire *et al*. 2014; Gardy and Loman 2018; Deng *et al*. 2020). They have the advantage of detecting as yet unidentified pathogens as well as avoiding “signature-erosion”, where mutations occur in primer targets resulting in false negative or positive results (Sozhamannan *et al*. 2015; Deng *et al*. 2020).

Serological evaluation is also important to map the geographical distribution of pathogens as well as assessing vaccine responses and community impact. Ebola virus (EBOV) serological surveys have been conducted more frequently due to the fact most of the human outbreaks are caused by EBOV (Mulangu *et al*. 2018; Brook *et al*. 2019). However, individuals with antibodies against Marburg virus have been detected in locations in West and Central Africa with no previous history of Marburg virus outbreaks (Steffen *et al*. 2020). In August 2021 the first case of MVD was confirmed in Guinea (WHO). After a rapid public health response, further cases have not been reported and in September 2021 the end of the outbreak was declared. Serological studies could be very useful should an unidentified disease outbreak occur, as unusual locations can hinder efforts to identify such outbreaks, such as in the large EBOV outbreak in west Africa. Serosurveillance of bats is equally important considering they are potential reservoirs for EBOV, raising the possibility of zoonotic spillover events (Nys *et al*. 2018; Laing *et al*. 2018). In the past decade, filovirus RNA has been detected in bats in Europe. These were designated as a new genus, *cuevavirus*, with a sole species, Lloviu (LLOV) virus, which recently re-emerged in bats in Hungary. Vesicular stomatitis virus (VSV) particles pseudotyped with the LLOV surface glycoprotein (GP) were shown to infect human cells *in vitro*(Negredo *et al*. 2011; Maruyama *et al*. 2014; Kemenesi *et al*. 2018; De Arellano *et al*. 2019); and recently isolation of infectious LLOV from an asymptomatic Schreiber’s bat was achieved. Monkey and human cell lines were permissive to LLOV (Kemenesi *et al*. 2022). Consequently, monitoring the distribution of different filovirus species in animals is important as they may pose the potential for a future spillover into humans.

Pseudotypes are chimeric non-replicative viruses encoding a reporter gene and bearing the glycoprotein of interest on its envelope. The use of pseudotyped virus particles has several advantages when studying highly pathogenic viruses as they can be handled in low-containment facilities, often yield high production titres permitting upscaled use, can be multiplexed for assaying different viruses and can be adapted for high-throughput screening. In addition, there is a range of reporter genes that can be incorporated, as well as being highly sensitive in neutralisation assays (Wright *et al*. 2008; Mather *et al*. 2013; Temperton, Wright and Scott 2015; Long *et al*. 2015; Ferrara and Temperton 2018; Scott *et al*. 2012) exhibiting strong correlation to the native study virus (Konduru *et al*. 2018). They have been used in tropism (Goldstein *et al*. 2018), vaccine evaluation (Ewer *et al*. 2016), antiviral screening (Xiao *et al*. 2018) studies, amongst others.

Most of the assays and methods described so far require high-power (−70/80°C) freezers for virus storage and expensive transportation requirements to maintain a cold chain to other laboratories or in-field facilities. Alternative methods, involving more modest temperature requirements for reagents would be advantageous for accessibility and cost. One solution to reduce those costs would be to use lyophilised reagents whenever possible, especially if these are to be sent to and used in tropical regions with high temperature and humidity. Lyophilisation or freeze-drying has been used in production of pharmaceutical products, such as vaccines, to avoid the need for cold chain transportation and to increase shelf life of reagents (Kraan *et al*. 2014). Lyophilisation usually consists of two steps: freezing of the sample followed by drying in a low-pressure environment, whereby frozen water in the sample sublimates in the first drying step (primary drying) and unfrozen water evaporates in the second drying step (secondary drying). The secondary drying step is performed at a higher temperature (20-40°C) to eliminate this moisture (Kraan *et al*. 2014; Nireesha *et al*. 2013; Wang 2000). For most of the current proof-of-concept study described here, drying was performed using a standard laboratory freeze-dryer, Up-scaling of the freeze-drying process was performed during a pilot proof of concept study using a pilot scale freeze-drier (Intravacc, Netherlands) to study whether product quality could be improved by drying at more controlled conditions.

Cryoprotectants are routinely added to samples prior to freeze drying in order to protect the integrity of the substance being lyophilised. Formulations suitable for freeze-drying are commonly prepared with sugars such as sucrose, trehalose and sorbitol dissolved in various buffer solutions (Wang 2000) and have been applied to the freeze drying of viruses, including recombinant adenoviruses and lentiviruses (Shin, Salvay and Shea 2010). We previously assessed the use of sucrose as a cryoprotectant for lyophilising pseudotyped viruses (PVs of influenza, rabies and Marburg viruses), followed by storage for up to one month at different temperatures and humidity conditions(Mather *et al*. 2014). PV titres were shown to be maintained in infectivity assays after resuspension of lyophilised pellets. Marburg virus PV titre recovery was near 100% after one-month storage at +20°C, as well as influenza and rabies PVs. In addition, reconstitution of the pellets with either cell culture medium or distilled water made no significant difference in these tests. Reconstituted influenza and rabies PVs continued to perform well in antibody neutralisation assays, where convalescent sera were available(Mather *et al*. 2014). In this study we aimed to assess stability (including air transport) of lyophilised filovirus PVs after significantly extended storage periods and, in addition to functionality in neutralisation assays.

## 2. Materials and Methods

### 2.1 Cells

CHO-K1 cells used as targets in neutralisation assays were a gift from Dr Emma Bentley and Dr Giada Mattiuzzo (NIBSC). HEK293T/17 cells were maintained in Dulbecco’s Modified Eagle Medium DMEM and CHO-K1 cells in Ham F12 Medium, both supplemented with 10% fetal bovine serum (FBS) and 1% penicillin/streptomycin (Pan Biotech) and kept at 37°C and 5% CO_2_.

### 2.2 Plasmids and pseudotype production

Pseudotypes used in this study were EBOV (Makona C15 Genebank accession number KJ660346), LLOV (Genebank accession number JF828358) and RAVV (Genebank accession number DQ447649). GP genes were encoded in the pCAGGS expression vector.

PVs were generated in T75 flasks (Thermo Scientific) by 3-plasmid transfection protocol: 1µg GP (300 ng GP for EBOV), 1.5µg pCSFLW luciferase reporter (Demaison *et al*. 2002), 1µg p8.91 HIV-1 gag-pol (Zufferey *et al*. 1997), using PEI (Sigma) at a ratio of 1:10 DNA:PEI in producer HEK293T/17 (ATCC CRL-11268) cells. Supernatants were harvested 48h post transfection and kept at −80°C.

### 2.3 Lyophilisation reagents and equipment

Sucrose (Sigma-Aldrich 84097-250G) was used as a cryoprotectant during lyophilisation. A stock solution was prepared to the desired final concentration in Dulbecco’s Phosphate-Buffered Saline (Pan Biotech).

Low surface-tension polypropylene 1.5 mL tubes (Simport, Canada T330-7LST) were used to prepare and lyophilise PV samples.

Unless indicated otherwise, lyophilisation was carried out in a FreeZone 2.5 L freeze-dryer (Labconco, USA), connected to a vacuum pump (Rotary Vane 7739402), except additional EBOV samples, which were prepared in Sucrose-DPBS cryoproctectant in the Viral Pseudotype (VPU) in Kent, frozen by placing into a −80oC freezer and shipped on dry ice to Intravacc (Bilthoven, The Netherlands) for lyophilisation in a Telstar Lyobeta freeze dryer. This allowed for the comparison between simple lab-based and pilot scale lyophilisation processes. Convalescent serum samples for antibody studies were obtained from the National Institute of Biological Standards and Control (NIBSC code 15/262).

### 2.4 Lyophilisation and storage of PVs

PV supernatant of known RLU/ml titre was mixed with 1M Sucrose-DPBS solution to a total volume of 200 µL at a 1:1 (v/v) ratio in a low-surface tension eppendorf tube, vortexed to mix contents, centrifuged briefly and placed at −80°C overnight. A needle-pierced lid was placed on top of each tube before freeze-drying to let vapour escape during pressure change in the lyophilisation process. The lyophilisation cycle was run overnight at −40°C to −50°C with pressure dropping to < 0.033 mBar (3.3 Pa).

After lyophilisation pierced lids were removed and the low-surface tension tubes’ own lids were closed before the freeze-dried samples were placed in experimental storage.

These storage conditions were: −20°C, +4°C, ambient temperature ∼+22.5°C, +37°C (20% humidity) and +37°C (90% humidity). Temperature and humidity were monitored regularly in the different storage containers with a Fisherbrand™ Traceable™ Jumbo Thermo-Humidity Meter (Fisher Scientific 11536973).

Reconstitution of lyophilised pellets was carried out in 100 µL of complete medium before titration or use in neutralisation assays.

For the lyophilisation performed at Intravacc, EBOV PV supernatant was mixed with 1M Sucrose-DPBS solution at a 1:1 (v/v) ratio, placed at −80oC overnight and shipped by air on dry ice, remaining frozen on receipt ∼24hrs later. EBOV samples lyophilised at VPU were also sent in the same shipment box, and stored at −80°C at Intravacc, then returned with the newly lyophilised samples to assess whether the journey had an impact on titre retention.

The materials used by Intravacc were as follows: Glass vials (APG Packaging 1003201), autoclaved in-house before use. Rubber stoppers (APG Packaging 1008739), in-house dried overnight at 105°C. Samples were thawed at RT. Forty glass vials were filled with 200 µl of PV supernatant and half-stoppered before loading in the Telstar Lyobeta freeze dryer. The sample vials were surrounded with empty vials in a metal fork.

After freeze-drying, the vials were fully stoppered in the freeze dryer, still under a pressure of 20 µbar. Subsequently, the vials were capped sealed with an aluminium cap.

A summary of the lyophilisation cycle is provided (Table 1) including the temperatures monitored during the cycle.

**Table 1.** Lyophilisation protocol performed at Intravacc.

| Freezing |  | Primary drying |  |  | Secondary drying |  |
| --- | --- | --- | --- | --- | --- | --- |
| Temperature (°C) | Time (hours) | Temperature (°C) | Time (hours) | Pressure (µbar) | Temperature (°C) | Time (hours) |
| -50 | - * | -45 | 0.5 | 20 | 25.5 | 24 |
| -50 | 2 | -45** | 96 | 20 | 25.5 | 24 |
|  |  | -45** | 2 | 20 | 4 | 0.5 |
|  |  |  |  |  | 4 | 99 *** |

\* Shelf preparation, prior to loading of the vials
\*\* Pressure rise test (PIM):
Max loops: 10
Extra drying time: 2 hours
Allowed pressure rise: 5 µbar
Test time: 60 seconds
\*\*\* Storage of the vials at 4°C until the freeze dryer was stopped manually

All lyophilised samples were transported back to the VPU by air. Lyophilised EBOV PV samples were then stored at −20°C, +4°C, +22.5°C, +37°C (20%) and +37°C (90%) for an initial 1-month period, and some for 6 months, at +22.5°C and +37°C (20%) to compare titre retention between lab and pilot scale lyophilisation processes.

### 2.5 Infectivity and neutralisation assays

Infectivity and neutralisation assays were performed as previously described (Wright *et al*. 2008; Temperton *et al*. 2007; Ferrara and Temperton 2018). Briefly, PV supernatant was added (100 µL/well) to a white, flat-bottom, sterile Nunc 96-well microplate (Thermo Scientific 10072151) and serially diluted 2-fold. Target cells (2 × 10^4^/well in 50 µL) were added and incubated for 48h at 37°C, 5% CO_2_. After 48h, the media was removed and discarded; Bright-Glo reagent (Promega) was added to the plate and incubated at room temperature for 5 min before measuring luminescence on a GloMax 96 luminometer (Promega).

In antibody neutralisation assays, a 2-fold serial dilution of serum was conducted in duplicate, in white, flat-bottom, sterile Nunc 96-well microplates, at a starting dilution of 1:40 and incubated with ∼100,000 RLU of PV at 37°C, 5% CO_2_ for 1h to allow the antibodies to bind to the GP. A PV only control (0% neutralisation equivalent) with no serum and cell only wells (100% neutralisation equivalent) was also set up. Target cells (2 × 10^4^/well) were added and incubated for 48h at 37°C, 5% CO_2_. After 48h, the plate was read as previously described.

The data was normalised to the percentage reduction in luminescence according to the average RLU of the cell only (100% neutralisation) and PV only (0% neutralisation) controls and fitted into a non-linear regression model (log [inhibitor] vs. normalised response – variable slope) to interpolate the inhibitory concentrations at 50% (IC_50_) and 90% (IC_90_), that is, the reciprocal of the dilution at which 50% or 90% of PV cell entry was inhibited, respectively.

Titre recovery of lyophilised PVs was calculated in comparison to their unlyophilised counterparts.

## 3. Results

### 3.1 PV production titres

Typical titres of ∼1 × 10^8^ RLU/mL were measured for EBOV and LLOV PV supernatants, and ∼1 × 10^10^ RLU/mL for RAVV PVs (Figure 1). Some of these PV supernatants were lyophilised and the remaining stock used as unlyophilised positive controls, as well as for comparison to calculate titre retention in further experiments.

**Figure 1.**
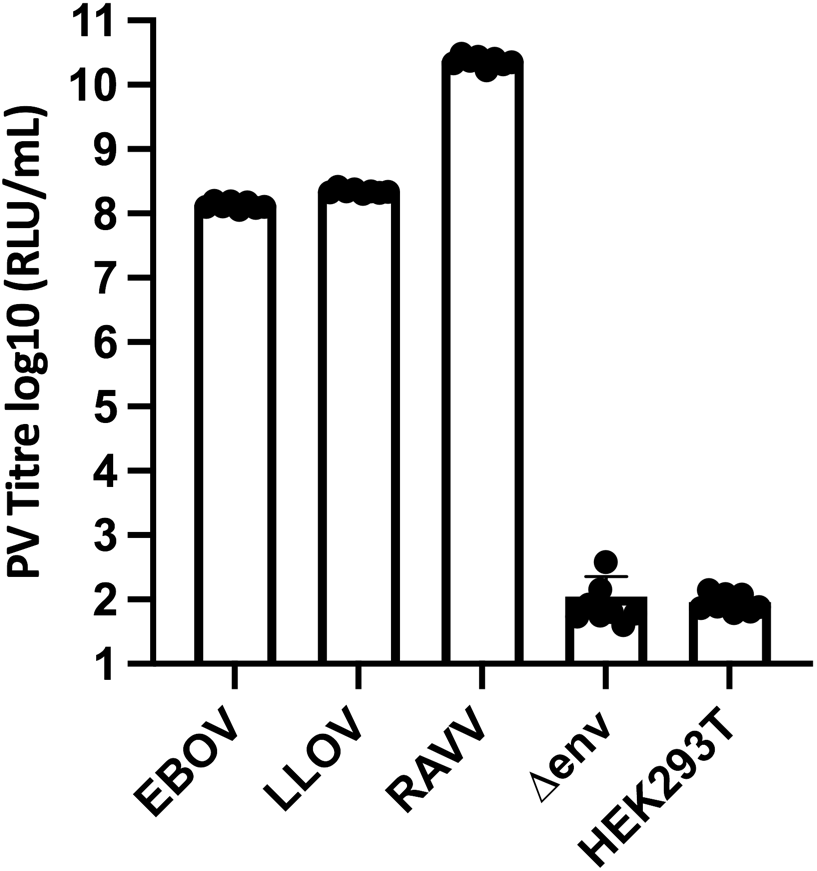
Generation of EBOV, LLOV and RAVV PVs. Transduction titres, as measured by PV-mediated luciferase reporter gene expression, are indicated as the mean ± s.d log_10_ (RLU/mL) values of at least three independent experiments. The titre of lentiviral particles bearing no GP (Δenv) and background luminescence from uninfected cells (HEK293T) is also shown.

### 3.2 Long-term storage of lyophilised PVs utilising a standard laboratory freeze-drier (Labconco)

Filovirus PVs lyophilised at the VPU earlier in the study were stored for up to 2 years under various conditions.

Impressively, EBOV PVs retained ∼100% of their titre when reconstituted after being stored at −20°C and +4°C for 2 years (Figure 2e). At higher temperatures, titres decreased to background levels within 6 months (Figure 2b). EBOV PVs retained 90% of their titre when reconstituted after being stored at +22.5°C for one month (Figure 2a), then titres decreased to background levels between one month and six months (Figure 2b). EBOV PVs stored at +37°C in dry (<25% humidity) and humid (90% humidity) conditions for one month did not generate any functional titres (Figure 2).

**Figure 2.**
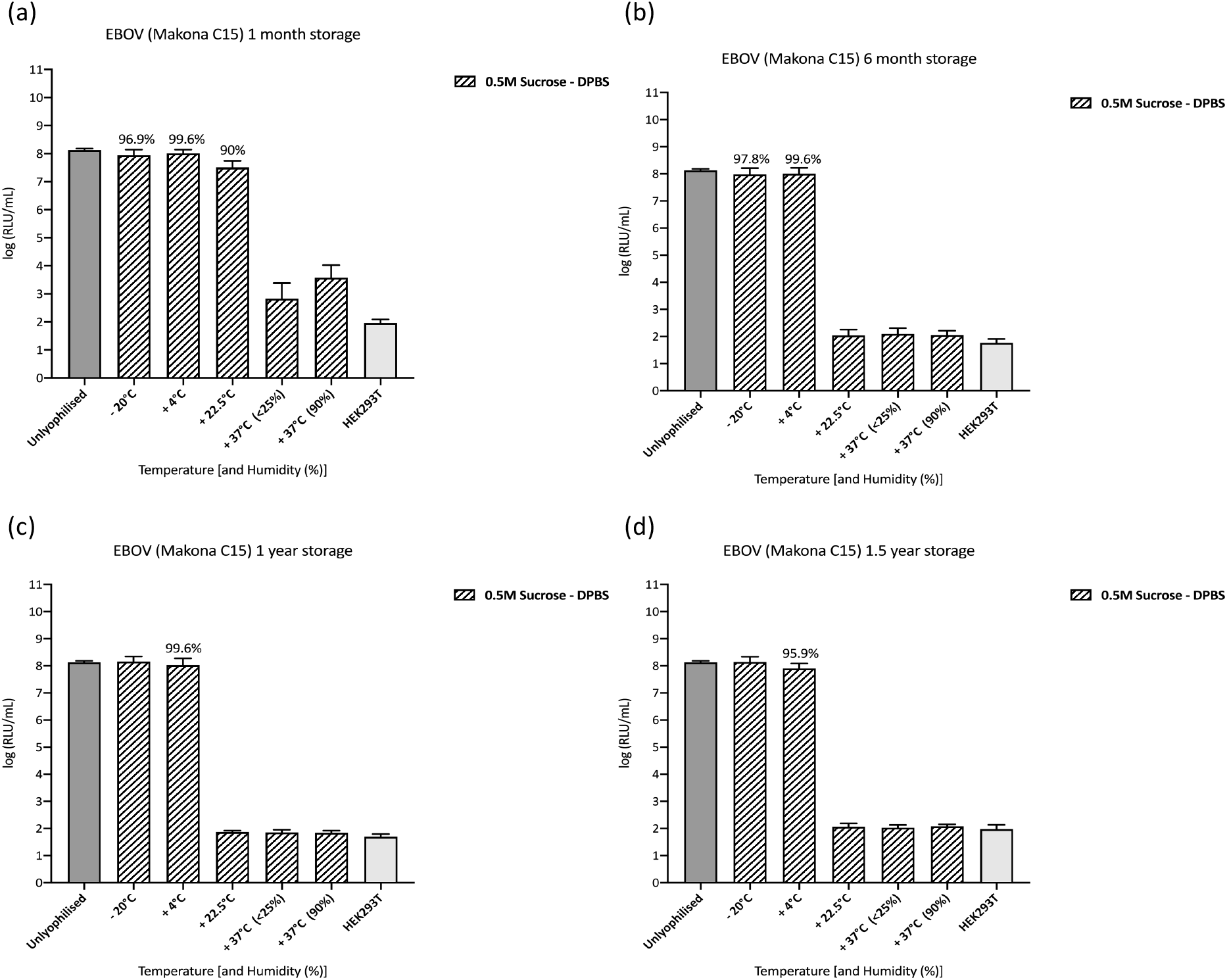
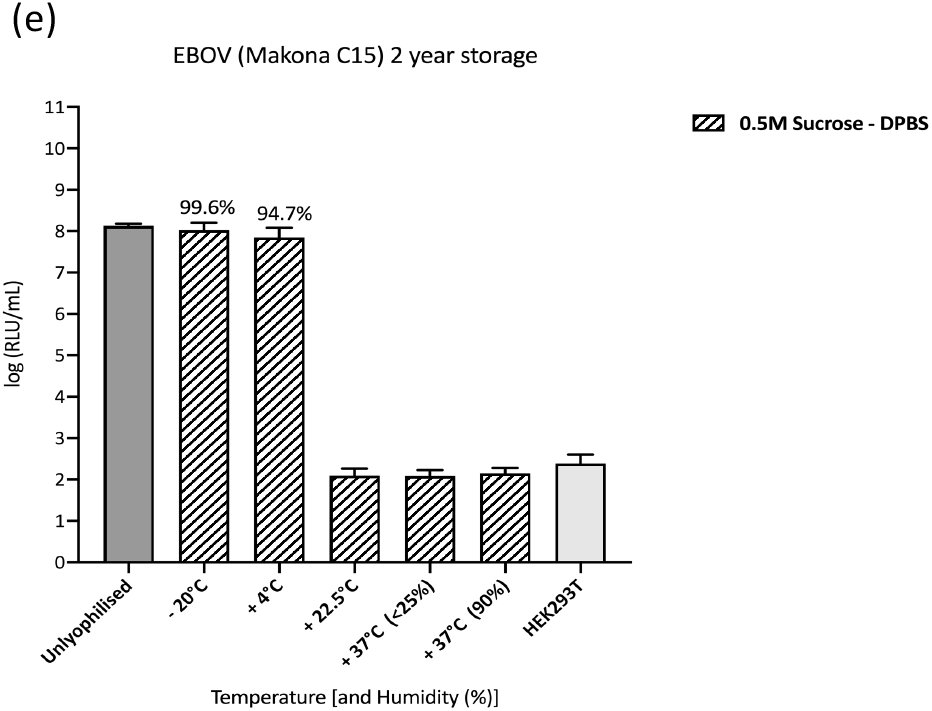
Infectivity assay following long-term storage of lyophilised EBOV PVs. Lyophilised PVs were stored at different temperatures and humidity conditions (<25% or 90%) for (a) 1 month, (b) 6 months, (c) 1 year, (d) 1.5 years and (e) 2 years. Unlyophilised EBOV PVs were positive controls for the assay, as well as a parameter for comparison to calculate titre retention after lyophilisation, storage and reconstitution. Transduction titres are expressed as the (log_10_) mean RLU/mL ± s.d from at least two independent experiments and % titre retention for functional titres are displayed on top of each bar if less than 100%, for those samples with measureable titre. Background luminescence in uninfected cells (HEK293T) is also shown.

For RAVV (Marburg virus) PVs, >90% of original titres were retained when reconstituted after being stored at −20°C and +4°C for up to 2 years (Figure 3). They retained 93.9% of their titre when reconstituted after being stored at +22.5°C for one month (Figure 3a), then titre recovery decreased to 69.5% between one month and six months (Figure 3b). At higher temperatures titres decreased to background level within 6 months (Figure 3b).

**Figure 3.**
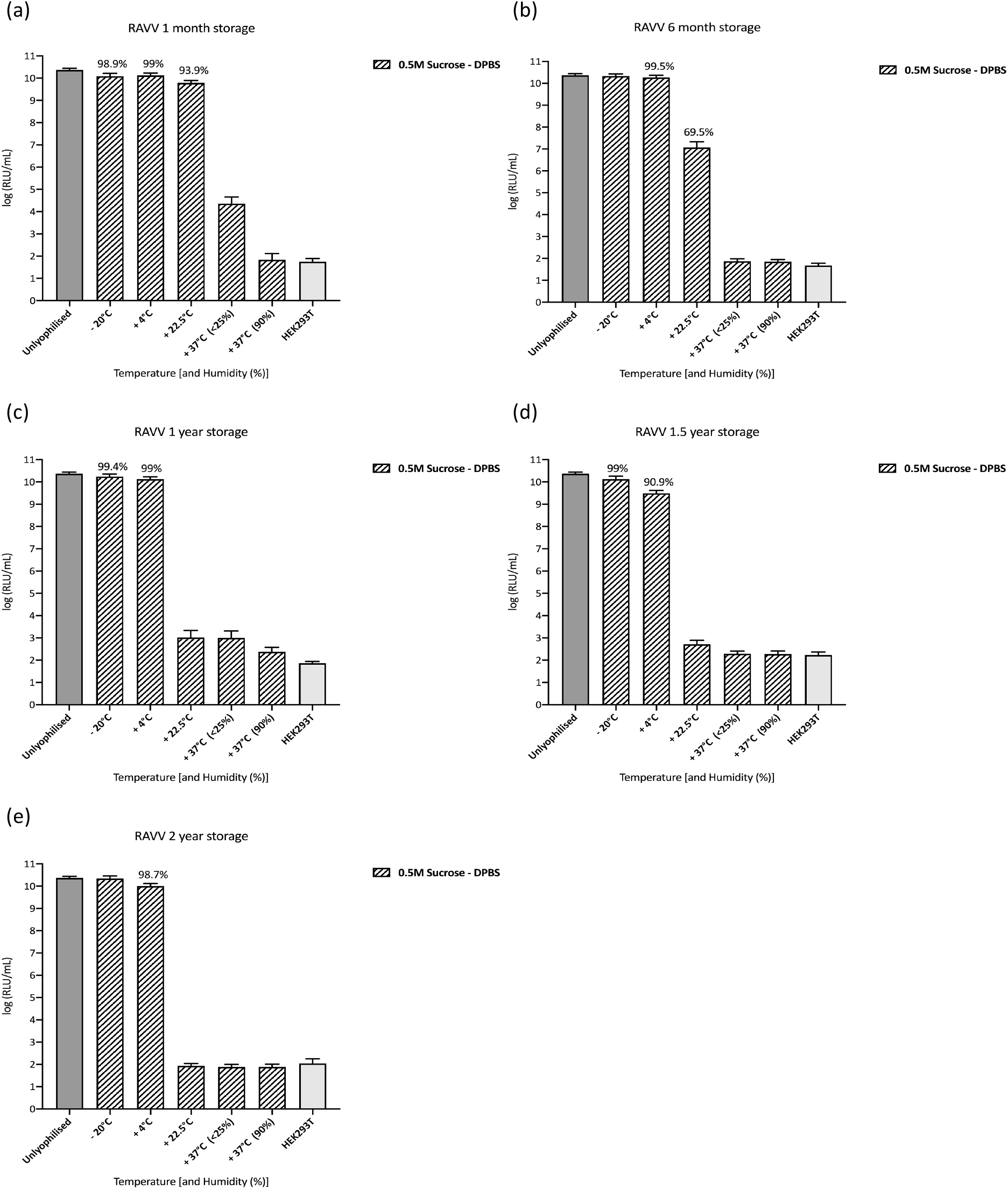
Infectivity assay following long-term storage of lyophilised RAVV PVs. Lyophilised PVs were stored at different temperatures and humidity conditions (<25% or 90%) for (a) 1 month, (b) 6 months, (c) 1 year, (d) 1.5 years and (e) 2 years. Unlyophilised RAVV PVs were positive controls for the assay, as well as a parameter for comparison to calculate titre retention after lyophilisation, storage and reconstitution. Transduction titres are expressed as the (log_10_) mean RLU/mL ± s.d from at least two independent experiments. Background luminescence in uninfected cells (HEK293T) is also shown.

RAVV PVs stored at +37°C in dry (<25% humidity) and humid (90% humidity) conditions did not generate any functional titres (Figure 3), although titres were slightly higher (∼10^4^ RLU/mL) than background level when stored for only one month at +37°C (<25% humidity) (Figure 3a).

Lastly, LLOV PVs (from the third filovirus genus) were lyophilised and stored for up to 1.5 years under different conditions.

LLOV PVs retained ∼90% of their titre when reconstituted after being stored at −20°C and +4°C for up to 1.5 years (Figure 4). At higher temperatures titres decreased to background level within 6 months (Figure 4b). LLOV PVs retained 84.9% of their titre when reconstituted after being stored at +22.5°C for one month (Figure 4a), then titres decreased to background level between one month and six months (Figure 4b).

**Figure 4.**
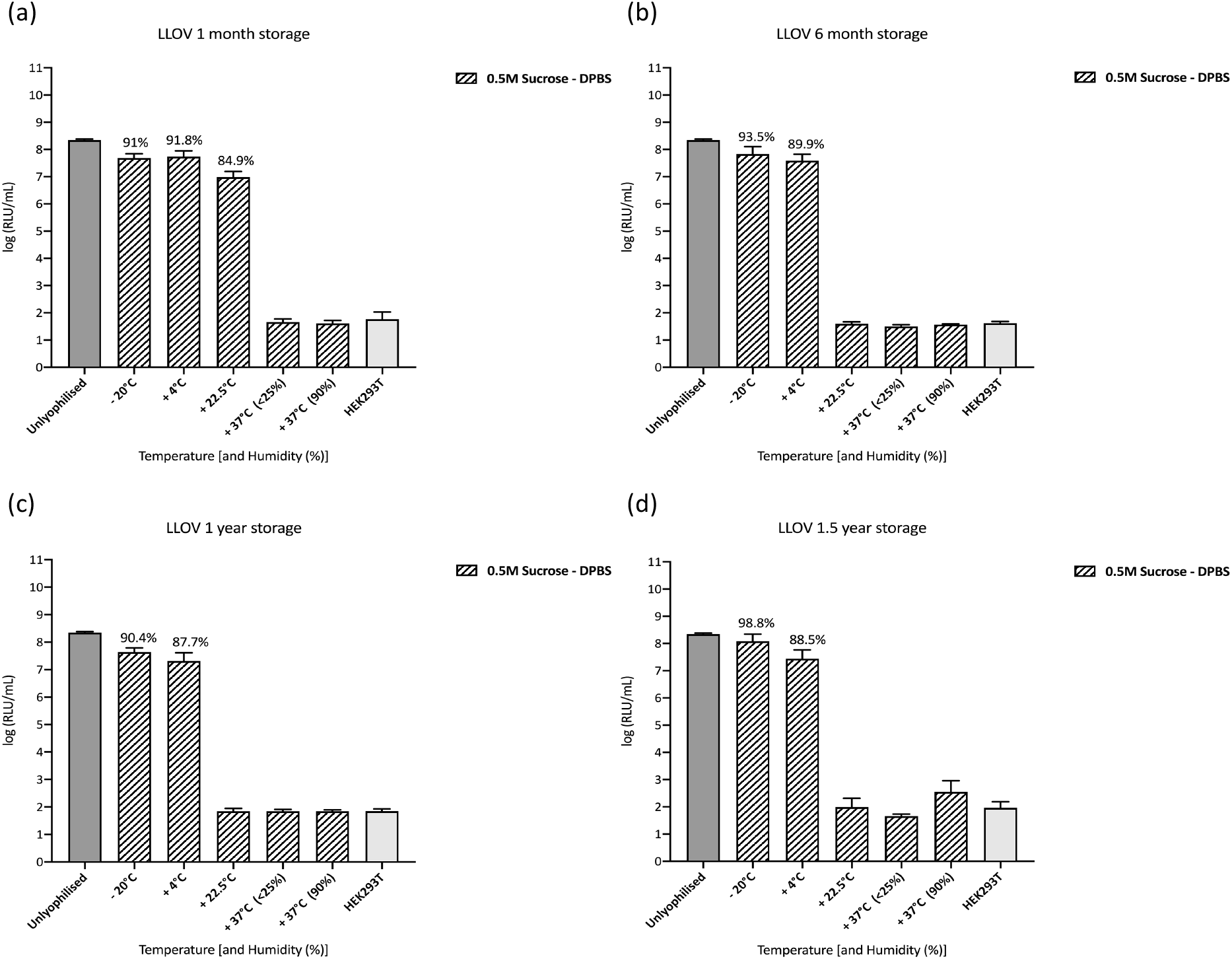
Infectivity assay following long-term storage of lyophilised LLOV PVs. Lyophilised PVs were stored at different temperatures and humidity conditions (<25% or 90%) for (a) 1 month, (b) 6 months, (c) 1 year, (d) 1.5 years. Unlyophilised LLOV PVs were positive controls for the assay, as well as a parameter for comparison to calculate titre retention after lyophilisation, storage and reconstitution. Transduction titres are expressed as the (log) mean RLU/mL ± s.d from at least two independent experiments. Background luminescence in uninfected cells (HEK293T) is also shown.

PVs stored at +37°C in dry (<25% humidity) and humid (90% humidity) conditions did not generate any functional titres after storage (Figure 4).

### 3.3 Short-term storage of lyophilised PVs comparing a standard laboratory freeze-drier Labconco) to pilot scale facilities (Intravacc)

A one-month storage stability assessment was performed. EBOV PVs lyophilised at the VPU that were shipped to The Netherlands, kept refrigerated for 2 weeks then shipped back to the VPU, retained titres after a further one month’s storage at −20°C and +22.5°C above 90%, however at higher temperatures titres were dropped to background levels (Figure 5a – blue). A temperature of +4°C was not tested, as retention at this temperature did not differ greatly from samples stored at −20°C in previous lyophilisation tests.

**Figure 5.**
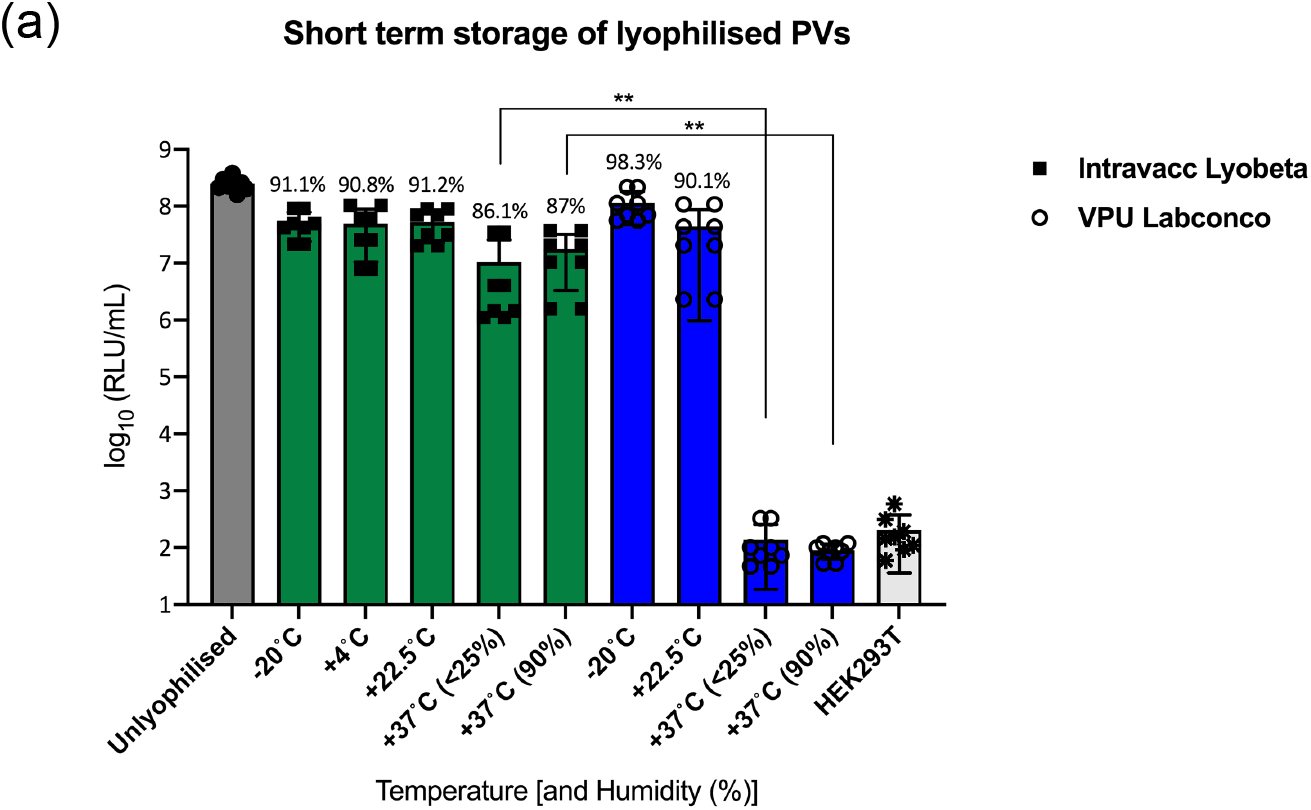
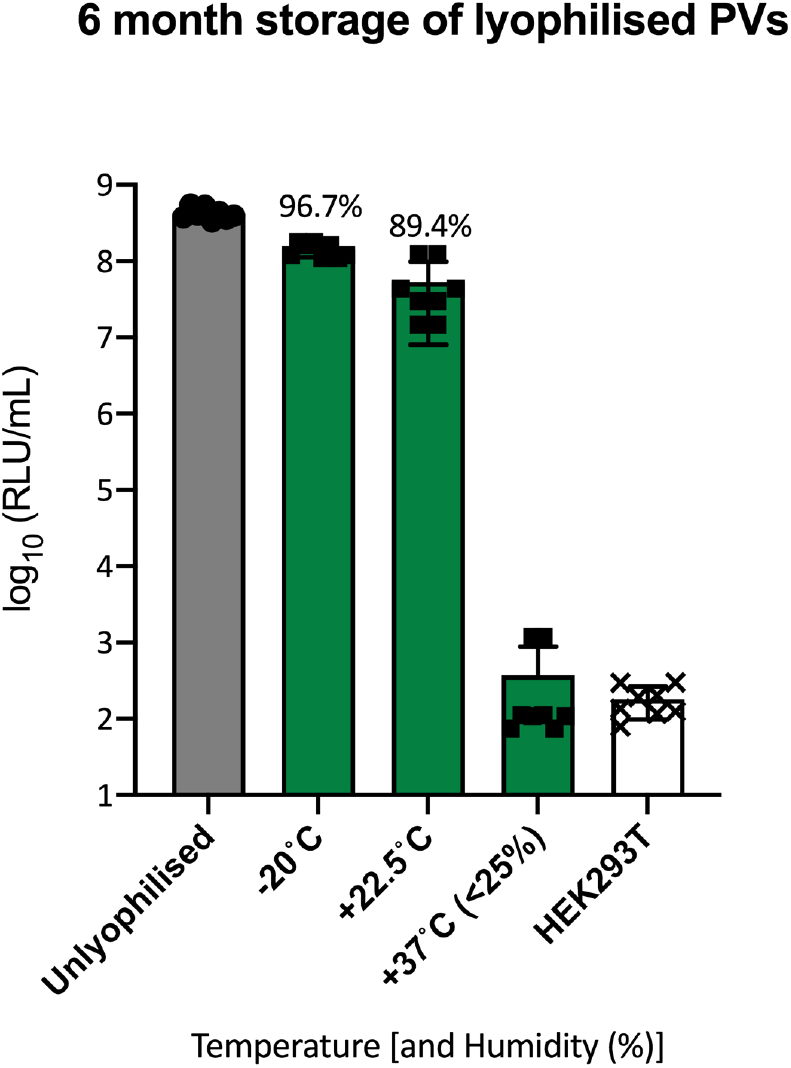
Infectivity assay following one-month and six-month storage of EBOV PVs lyophilised at Intravacc. PVs were lyophilised at Intravacc (green) or at the Viral Pseudotype Unit (VPU; blue) and stored at different temperatures and humidity conditions (<25% or 90%) for (a) 1 month and (b) 6 months. Unlyophilised EBOV PVs were employed as positive controls for the test, as well as a parameter for comparison to calculate titre retention after lyophilisation, storage and reconstitution. Titre recovery %) is expressed on top of each bar. **p<0.01 (Mann-Whitney test). Background luminescence in uninfected cells (HEK293T) is also shown.

By contrast, EBOV PV samples lyophilised at Intravacc withstood storage at high temperatures of +37°C in dry or humid conditions for at least a month (Figure 5a – green), with 86.1% of titre recovered after being stored at +37°C (<25% humidity) and 87% of titre recovered after being stored at +37°C (90% humidity). This represents a significant increase (p<0.01) in stability and recovery when compared to the EBOV PV samples lyophilised in the VPU Labconco freeze-dryer (Figure 5a).

Samples lyophilised at Intravacc stored at −20°C, ambient temperature (+22.5°C) and +37°C (<25% humidity) were further assessed after 6 months (Figure 5b). For PV samples stored at lower temperatures (−20°C and +22.5°C) titre retention was above 89.4%, however for samples stored at +37°C (<25% humidity) PV titres had decreased to background (HEK293T) levels (Figure 5b).

### 3.4 Pseudotype neutralisation assay with lyophilised samples

Following demonstration that lyophilised, stored and reconstituted PVs retained ability to transduce target cells, EBOV samples were then tested for biological functionality in antibody neutralisation assays.

Lyophilised EBOV PVs in Intravacc’s Telstar Lyobeta freeze dryer stored at −20°C, +37°C (<25%) and +37°C (90%) were reconstituted to be used as input in PVNAs against pooled convalescent EBOV serum (WHO standard NIBSC 15/262) to compare performance with unlyophilised PVs (Figure 6a). EBOV PVs lyophilised in the Labconco freeze-drier that had been stored for 1.5 years at +4°C then reconstituted and used as the PV input in PVNA (Figure 6b).

**Figure 6.**
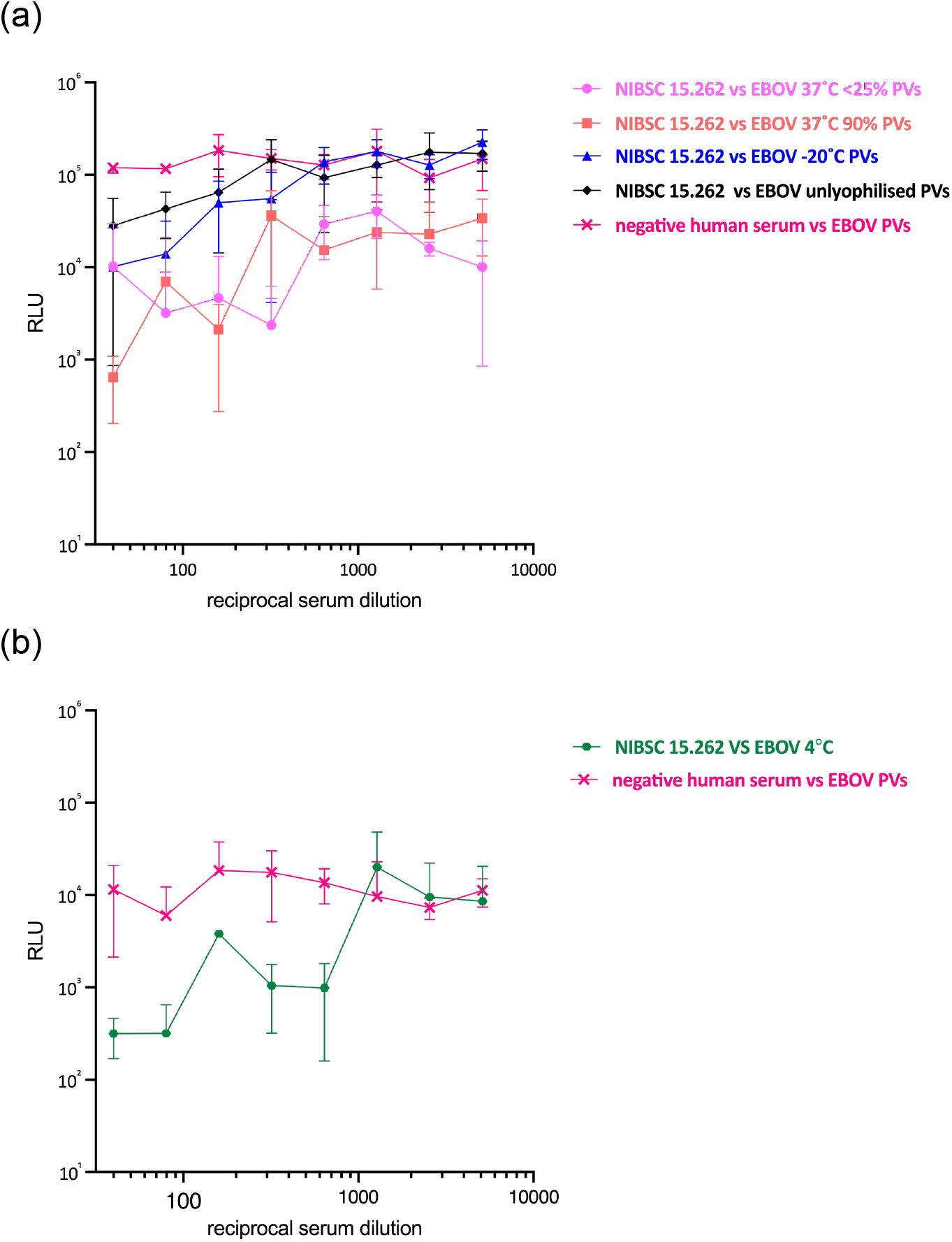
Neutralisation assay using reconstituted lyophilised EBOV PVs. WHO standard NIBSC 15.262 tested against EBOV PVs lyophilised with: (a) Telstar Lyobeta that had been stored for 1 month at −20°C, +37°C (<25% humidity), +37C (90% humidity); and (b) Labconco stored for 1.5 years at +4°C. Negative human serum from a healthy donor (Sigma) was used as a negative control. Decrease in target cell luminescence (mean ± s.d) from duplicates in two independent experiments,

Despite the neutralising response being variable between samples, especially those that had been stored at higher temperatures (Figure 6a), they all were able to detect neutralising antibodies in the convalescent test serum as evidenced by the reduction in reporter gene expression, with PVs stored at −20°C more comparable IC_50_ to unlyophilised PVs (Figure 6, Table 2).

**Table 2.**
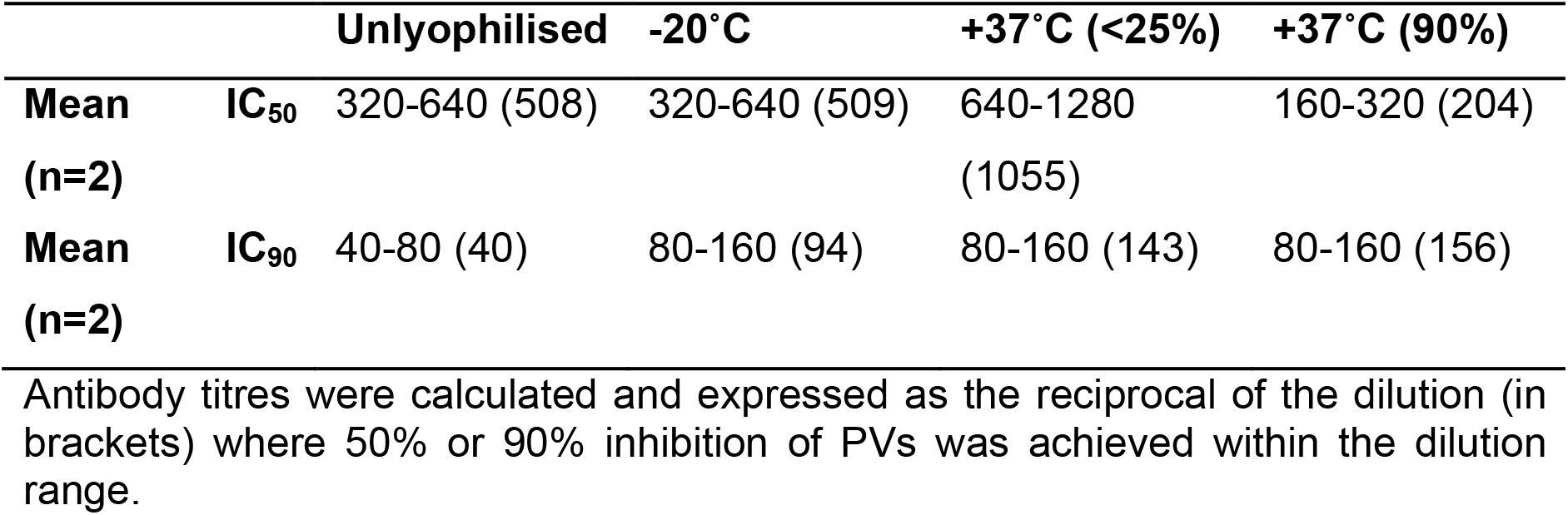
Mean half-maximum inhibitory concentration (IC_50_) and 90% inhibitory concentration (IC_90_) of Intravacc lyophilised samples in PVNAs.

## 4. Discussion

The need to improve diagnostics and serological tests for emerging diseases was illustrated during the large EBOV outbreak in West Africa in 2013-2016 (Murphy 2019; Gatherer 2014; Formella and Gatherer 2016), and more recently during EBOV outbreaks in the DRC and Guinea with over 50 people affected as well as the first case of MARV reported in West Africa in 2021 (Centers for Disease Control and Prevention). Other emerging diseases such as measles in the DRC, concurrent with EBOV, and therefore increasing the burden on health services as well as the ongoing SARS-CoV-2 outbreak with over 500 million confirmed cases globally (WHO-12/05/2022) have also stressed the need for research in emerging diseases.

Although RT-PCR based assays are the gold standard for diagnostic testing of filoviruses and other viruses such as SARS-CoV-2 (Osterdahl *et al*. 2020; Clark *et al*. 2018), these require the presence of virus genetic material in patient blood or tissues. Serology looks for the antibodies in humans and animals to reveal the imprint of infection, even in individuals that did not exhibit clinical signs. Seroepidemiological studies can reveal geographical distribution, zoonotic spillover and can be used retrospectively to detect historical infections (Mather *et al*. 2013; Ewer *et al*. 2016; Kinsley, Scott and Daly 2016; Luczkowiak *et al*. 2016).. A major issue with conducting neutralising antibody studies however is the use of native virus, and the need to handle potentially dangerous pathogens in high containment facilities.. This particularly impacts the ability for field laboratory work. The application of pseudotype viruses (PVs) to such studies may provide a suitable alternative, as long as the essential reagents can be provided in a stable form. A PV based neutralising antibody assay that could differentiate between genera and even species of filoviruses and a complementary ELISA would be highly desirable to provide epidemiological data, as well as monitoring outbreaks when cross-reactivity might be an issue and next generation sequencing is not available. Importantly, PVs have the advantage of only requiring low containment facilities (BSL 1-2), and being amenable to multiplexing (Carnell *et al*. 2015; Ewer *et al*. 2014). Like native virus neutralisation assays, PVNAs require a suitable target cell line exhibiting the virus-specific receptor, but only need one to two days to obtain results

The majority of emerging virus outbreaks, such as those caused by filoviruses has occurred in low-resource countries. In order to provide assay reagents to laboratories in these regions there is often a need for cold-store transportation and storage to prevent deterioration. Lyophilisation has been utilized as a means to address this issue and permit the transport of certain reagents and vaccines in ambient conditions (Kraan *et al*. 2014; Bjelošević *et al*. 2018; Bjelošević *et al*. 2020; Wong, Al-Salami and Dass 2020).

The possibility of using lyophilisation for short-term storage of PVs for antibody detection was explored for influenza, rabies and Marburg viruses (Mather *et al*. 2014).Generally, PVs retained titres after storage at lower temperatures of −80°C up to room temperature (20°C) when stored for up to one month. However, at +37°C in dry or humid conditions, PV titres decreased approximately 100-fold, though still retaining functionality due to their initial high titre. Reconstituted lyophilised influenza and rabies PVs also continued to perform in PVNAs. Marburg PVs were not tested in PVNAs due to lack of available anti-sera.

In the current study, we undertook a more comprehensive analysis of the application of lyophilisation to PVs with subsequent stability and functionality testing. This involved investigating different lyophilisation equipment and protocols, long-term storage under a range of conditions. Reconstituted PVs were tested in titration and neutralisation assays. We compared a standard laboratory freeze-dryer (Labconco) to pilot scale freeze-dryer (at Intravacc, The Netherlands). The disaccharide sucrose was used as a cryoprotectant for freeze-drying as successful use had been established previously (Nireesha *et al*. 2013; Kraan *et al*. 2014; Mather *et al*. 2014).

All PVs were assessed in infectivity assays to calculate titre retention. The availability of convalescent patient antisera enabled the functionality of lyophilised and non-lyophilised EBOV PVs to be compared in neutralisation tests (n.b. no specific antisera was available for other filovirus genera).

All PVs utilised in this study had a lentiviral core. Initial PV titres for *ebolaviru*s (EBOV) and *cuevavirus* (LLOV) PVs were 1 × 10^8^ RLU/mL, whereas for *marburgvirus* (RAVV) PVs was 1 × 10^10^ RLU/mL (Figure 1), consistent with those generated in our previous study (Mather *et al*. 2014).

Following production, PVs were then lyophilised using a standard laboratory freeze dryer, placed under a range of storage conditions then sampled and titrated at various time intervals over a two year period. At higher temperatures, we observed a large drop in transduction titres after only one-month storage at +37°C in dry and humid conditions (Figure 2-4a).

For assessment of long-term storage and stability of lyophilised samples, PVs were generated then mixed with Sucrose-DPBS cryoprotectant solution to a final concentration of 0.5M before freeze drying. All three genera of lyophilised filovirus PVs followed a similar trend after long-term storage.

All lyophilised PVs had titre recovery above 85.9% when stored in a household fridge at +4°C for 1.5 years (Figure 2-4). Furthermore, EBOV (Figure 2e) and RAVV (Figure 3e) PVs had titre recoveries of 94.7% and 98.7% respectively after being stored at +4°C for 2 years. These are particularly encouraging results as avoiding the need for high-powered freezers would expand the number of labs in low-resource countries, such as those involved in the recent filovirus outbreaks, being able to employ PVs for research and assays,

For samples stored at ambient temperature (+22.5°C), the decrease in titre recovery between one and six months was not investigated further, assuming that fridge storage could be achieved within a month, following transport and delivery.

We hypothesized that the rapid decrease in titres at 37°C could be due to the residual moisture remaining after lyophilisation without employing further pellet drying (Nireesha *et al*. 2013). Consequently we sent frozen EBOV PV samples in sucrose/DPBS (final concentration 0.5M; on dry ice) to be lyophilised using a pilot scale freeze-dryer at Intravacc including a secondary drying step. Samples were then transported back (on dry ice) for analysis at the VPU (Figure 7). EBOV PVs lyophilised at Intravacc had titre recovery of 86.1% and 87% even after being stored for a month at +37°C in dry and humid conditions (Figure 5a-green). The samples for treatment at Intravacc were accompanied by a set that had been lyophilised at the VPU (see data in Figure 5a – blue), which were temporarily stored at −80°C in the Netherlands then returned to the VPU on dry ice. By contrast, these samples had a drastic drop in titres after a month’s storage at +37°C. Samples lyophilised at Intravacc and then stored at +37°C (<25% humidity) only began to lose titre after between one and six months of storage (Figure 5). Overall, these are very encouraging results as PVs lyophilised using pilot scale equipment will retain a functional titre even when stored at harsher, warmer conditions. This suggests lyophilised PVs could be transported at room temperature to warmer tropical countries.

**Figure 7.**
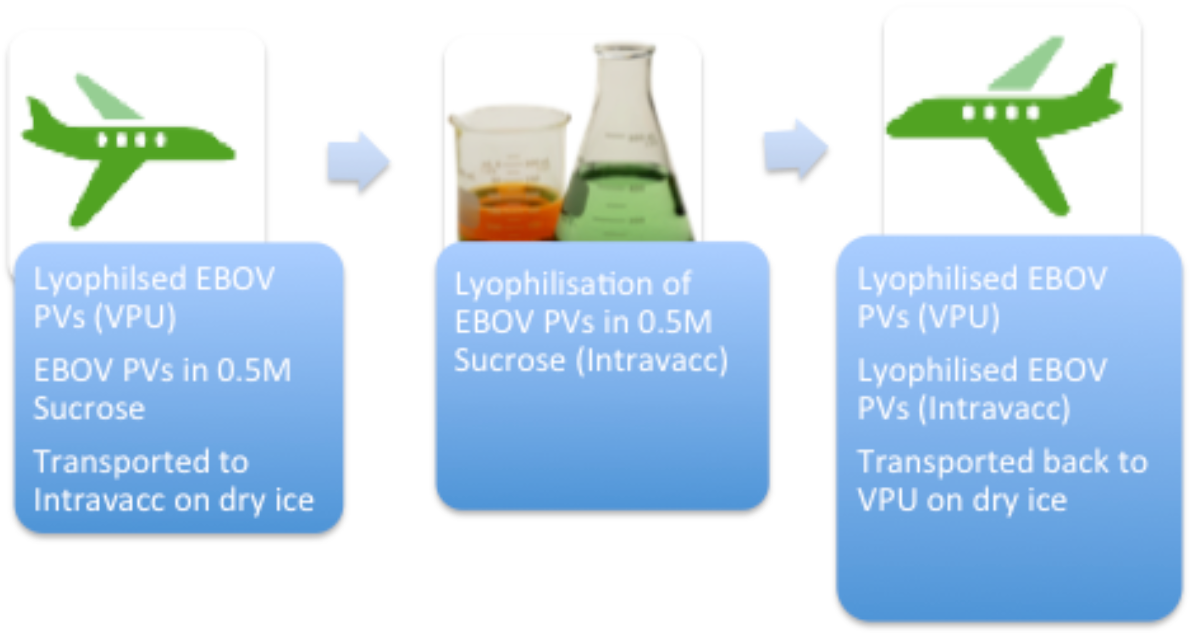
Diagram showing the transportation of lyophilised PVs between Kent (UK) and Bilthoven (NL)

Finally, to assess performance of lyophilised PVs in PVNAs, EBOV samples lyophilised at the VPU that had been stored at +4°C for 1.5 years were tested and performed to standard in PVNAs (Figure 6b). EBOV PVs lyophilised at Intravacc also performed to standard in PVNAs, including samples stored at +37°C for one month (Figure 6a). They all detected neutralising antibodies in the convalescent serum, even if with somewhat variable responses (Table 2).

Other filovirus genera were not assessed in PVNAs in this study due to the lack of specific convalescent sera, however, they could be tested in PVNAs against monoclonal antibodies with neutralising activity in the future, as proof-of-principle. It would also be useful to assess performance of filovirus VSV core PVs in lyophilisation studies as VSV is widely used as a PV core for filoviruses(Takada *et al*. 1997; Maruyama *et al*. 2014; Ilinykh *et al*. 2016; Ruedas *et al*. 2017; Salata *et al*. 2019).

Overall, these results are very promising for a future serological kit that could be transported at ambient temperature and would last at least two years in a household fridge. The lyophilisation of mammalian cells has been explored but so far has proved elusive, although there has been some success in lyophilisation and reconstitution of platelets (Wolkers, Tablin and Crowe 2002), or somatic cells that have been lyophilised then used in nuclear transfer experiments (Loi *et al*. 2008). However, lyophilising mammalian cells for later propagation in culture has not been successful so far due to the integrity of the cell membrane being compromised and the resulting damage (Zhang *et al*. 2017). In that case, cells would have to be sent as a frozen stock for propagation.

## 5. Conclusion

This data demonstrates that PVs lyophilised using either lab-based or pilot scale systems are functional in antibody neutralisation assays, after short-term storage in tropical conditions or after several years in standard refrigeration. This important finding would permit the widespread employment of PV-based antibody assays in countries that have historically suffered from filovirus outbreaks.

## Acknowledgements

We would like to thank Intravacc for their research support.

## Notes

### Competing Interest Statement

The authors have declared no competing interest.

